# Adaptive significance of quorum sensing-dependent regulation of public goods by integration of growth rate: a trade-off between survival and efficiency

**DOI:** 10.1101/042739

**Authors:** Arvin Nickzad, Eric Déziel

## Abstract

Quorum sensing (QS) is a mechanism of communication used by bacteria to monitor cell density and coordinate cooperative behaviors. An emerging framework is that the adaptive significance of QS in regulation of production of costly extracellular metabolites (public goods) is to maintain the homeostasis of cooperation. We investigated the functionality of QS-dependent regulation of rhamnolipids, extracellular surface-active glycolipids promoting the social swarming motility behavior, in *Burkholderia glumae* and found that QS is superfluous under rich nutritional conditions. In contrast, decreasing nutrient concentrations to reduce the growth rate amplifies rhamnolipid biosynthesis gene expression, revealing a system where QS-dependent regulation is triggered by the growth rate of the population rather than by its cell density. Our results provide evidence that the adaptive significance of QS in regulation of public goods is to maintain an optimized demand-driven supply of target cooperative behavior, wherein efficiency can be traded off against survival.

## Introduction

Quorum sensing (QS) is a cell density-dependent mechanism which enables a group of bacteria to synchronize the expression of behaviors in response to the accumulation of self-produced autoinducer signals in their local environment (Fuqua et al., 1994). Such auto-induction regulatory systems are implicated in regulation of various cooperative phenotypes ranging from bioluminescence, virulence and nutrient acquisition to multicellular swarming motility and biofilm development (Passador et al., 1993, Bainton et al., 1992, Stevens and Greenberg, 1997, Daniels et al., 2004, An et al., 2014, Davies et al., 1998). These behaviors are considered cooperative because they mostly involve the production of costly extracellular metabolites («public goods») which, once produced, are available to all members of the group to share their benefits (Platt and Bever, 2009). Examples among bacteria are widespread and include production of siderophores, exoenzymes or surfactants (West et al., 2006). While cooperative production of public goods enhances the fitness of the whole population, at the individual level it is prone to exploitation by non-cooperative individual cells («cheaters») that, by not producing the metabolite or QS signal, gain a competitive edge over cooperative cells and hence threaten the homeostasis of the cooperation (Sandoz et al., 2007, Diggle et al., 2007); this brings up questions about the evolutionary stability of cooperation.

In recent years, the adaptive functionality of QS in regulation of public goods has constituted the main subject of several studies, and has been reviewed extensively (Schuster et al., 2013, Hense and Schuster, 2015). The central question relates to the benefits provided by QS in the production of public goods, and thus its role in the evolution and stability of cooperative behaviors (Schuster et al., 2013). In this regard, few studies addressing the social function of QS in regulation of public goods have demonstrated that QS can offer density-dependent fitness benefits (Pai et al., 2012, Darch et al., 2012, Goo et al., 2012). QS is used to coordinate the triggering of social behaviors at high cell densities when they are more efficient and will provide the greatest advantage to the community (Darch et al., 2012). Moreover, the social function of QS in regulation of public goods is not only limited to providing fitness benefits at higher cell densities, as it can also serve as a means to anticipate stationary phase or foresee the optimum cell density of a population (Goo et al., 2012). Cooperation presents a social dilemma, as cheaters should have a fitness advantage over cooperators. Still, QS can impose metabolic constraints on social cheating and protect the population from a ‘tragedy of the commons’ by simultaneously controlling the expression of private goods (Dandekar et al., 2012). Another possible solution to stabilizing cooperation is policing, where cooperators actively penalize social cheats to hinder their success (Wang et al., 2015).

While these studies offer insights on the role of QS in the evolution and stability of cooperative behaviors, the adaptive significance of QS has remained debatable. In fact, it was suggested that it is only through studying each specific auto-induction system in its evolutionary and ecological context that we can figure out its adaptive significance (Platt and Fuqua, 2010). The general framework is that, while conditions specific to each ecological context, such as population cell density and spatial diffusion properties, determine whether triggering of the costly QS-regulated cooperative behaviors is beneficial to the population, the integration of QS circuitry with other global regulatory pathways such as stress responses, specific environmental and physiological cues ensures an optimized demand-driven supply of target cooperative behaviors (Hense and Schuster, 2015). Indeed, integration of metabolic information into decision-making processes governing production of public goods, called ‘metabolic prudence’, can stabilize cooperative behaviors and reduce the cost of cooperation for the individual cell (Boyle et al., 2015, Xavier et al., 2011). Thus, the emerging picture is that in order for QS to properly regulate costly bacterial cooperative behaviors (when benefits outweigh costs of production) it must be integrated with additional physiological cues (Hense and Schuster, 2015).

Here, we demonstrate that QS-dependent regulation of a public good in the Gram-negative bacterium *Burkholderia glumae*, a pathogen of rice, is dependent on growth rate, a direct measure of the carrying capacity of a population through prevailing environmental conditions. *B. glumae* uses the LuxI/LuxR-type QS system TofR/TofI to regulate protein secretion, oxalate production and expression of various virulence determinants such as toxoflavin, lipase, KatG and flagella (Kim et al., 2007, Goo et al., 2010, Goo et al., 2012, Chun et al., 2009). The TofR QS transcriptional regulator is activated by its cognate *N*-octanoyl homoserine lactone (C_8_-HSL) autoinducer, the product of the TofI *N*-acyl-HSL synthase, to control the expression of target genes (Kim et al., 2004). Similarly to what is seen in *Pseudomonas aeruginosa* (Ochsner and Reiser, 1995, Köhler et al., 2000), we have previously demonstrated that in *B. glumae* as well, QS positively regulates the expression of the *rhlA* gene, encoding the enzyme catalyzing the first step in biosynthesis of rhamnolipids. These extracellular surface-active glycolipids promote the social surface behavior called swarming motility (Nickzad et al., 2015). Now, we show that, in contrast to *P. aeruginosa*, this QS regulation is dispensable in *B. glumae*, as rhamnolipid production is unaffected in a QS-negative mutant under unrestricted, rich nutritional conditions. This unexpected observation led us to investigate the interplay between QS and nutrition-based regulation of rhamnolipids in *B. glumae*. We found that growth rate, and not cell density itself, is the key factor triggering the QS-dependent regulation of rhamnolipid biosynthesis. Since rhamnolipids act as public goods promoting social swarming motility, integration of growth rate with QS-dependent regulation of rhamnolipids in *B. glumae* supports the concept that the adaptive significance of QS lies in providing a regulatory mechanism for trade-off between survival and efficiency.

## Results

### Quorum sensing regulation of rhamnolipid production in *B. glumae* is nutritionally conditional

To strengthen our understanding on the specific role of QS regulation in the production of rhamnolipids in *B. glumae* we studied the kinetics of rhamnolipid production in wild-type strain and *toƒl^-^* mutant using a minimal (mineral salts medium; MSM) typically used to promote production of rhamnolipids and a complex (nutrient broth; NB) culture media, both supplemented with non-limiting glycerol as carbon source (thus MSMG and NBG). As expected, the *toƒl^-^* mutant that lacks the production of C_8_-HSL does not produce rhamnolipids in the minimal medium and this production is restored by adding C_8_-HSL (Fig. 1a). On the other hand, unexpectedly, the *toƒl^-^* mutant presents no defect in production of rhamnolipids in rich NB medium (Fig. 1b).

**Figure 1.**
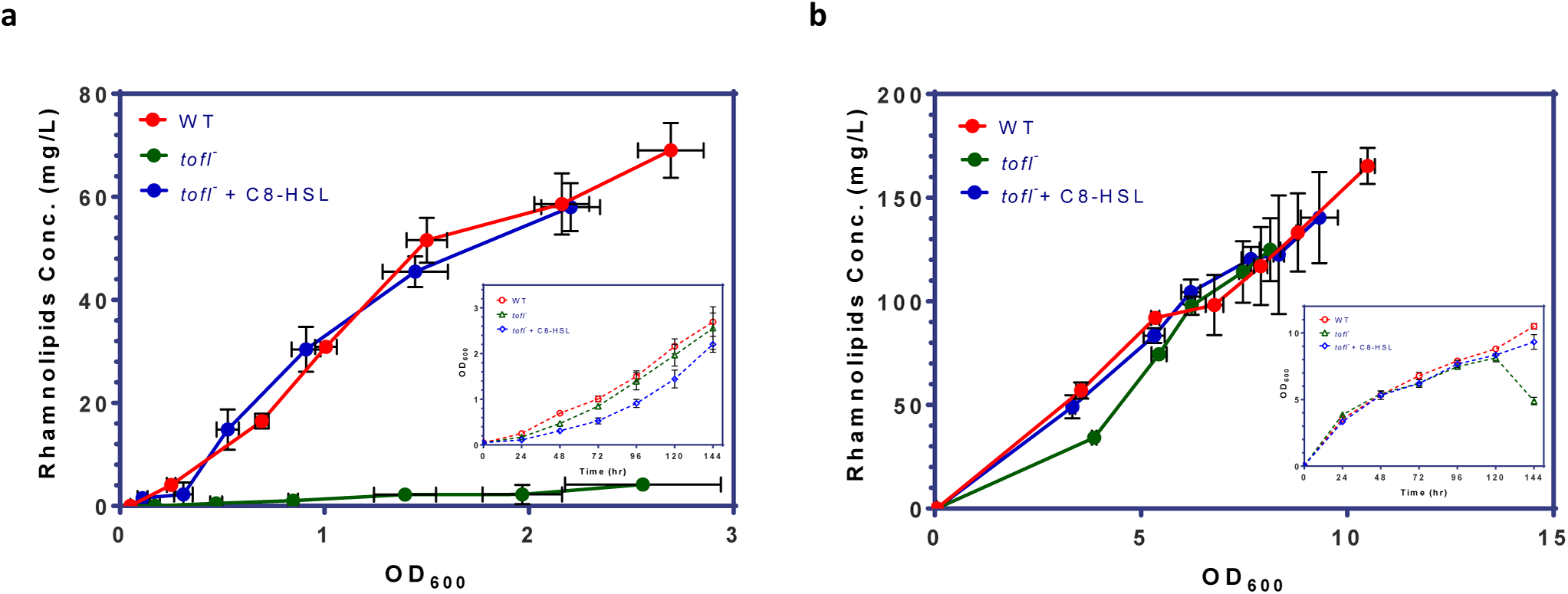
Rhamnolipid production of *B. glumae* strains in minimal vs. complex media. (a) While the wild-type strain produces rhamnolipids during culture in MSMG medium, the *toƒl^-^* mutant is not capable of producing rhamnolipids. The wild-type production level is restored upon exogenous addition of 8 μM C_8_-HSL to *toƒl^-^* mutant culture medium. (b) In contrast, in NBG medium the *toƒl^-^* mutant is capable of producing rhamnolipids at wild-type levels. The error bars indicate the standard deviation of the mean for three independent cultures. Insert graphs show the growth data plotted against time for the all the strains.

### Quorum sensing-dependent regulation of rhamnolipids is coupled to growth rate

Since a key difference between the two culture conditions is slower growth in MSMG vs NBG (Fig. 1), we hypothesized the impact of QS on biosynthesis of rhamnolipids is influenced by the nutritional conditions. We thus compared the rhamnolipid yield per biomass in wild-type and *toƒl^-^* mutant in increasing dilutions of NB medium, while maintaining the glycerol at 4%, in order to have a decreasing gradient of nutrients.

By calculating the ratio of specific production yields of the wild-type over the *toƒl^-^* mutant, we found that the contribution of QS on rhamnolipid production rises linearly with increasing NB medium dilution (Fig. 2). This shows that the regulation of rhamnolipid production relies more and more on QS upon increasing the nutrient limitation, supporting the hypothesis.

**Figure 2.**
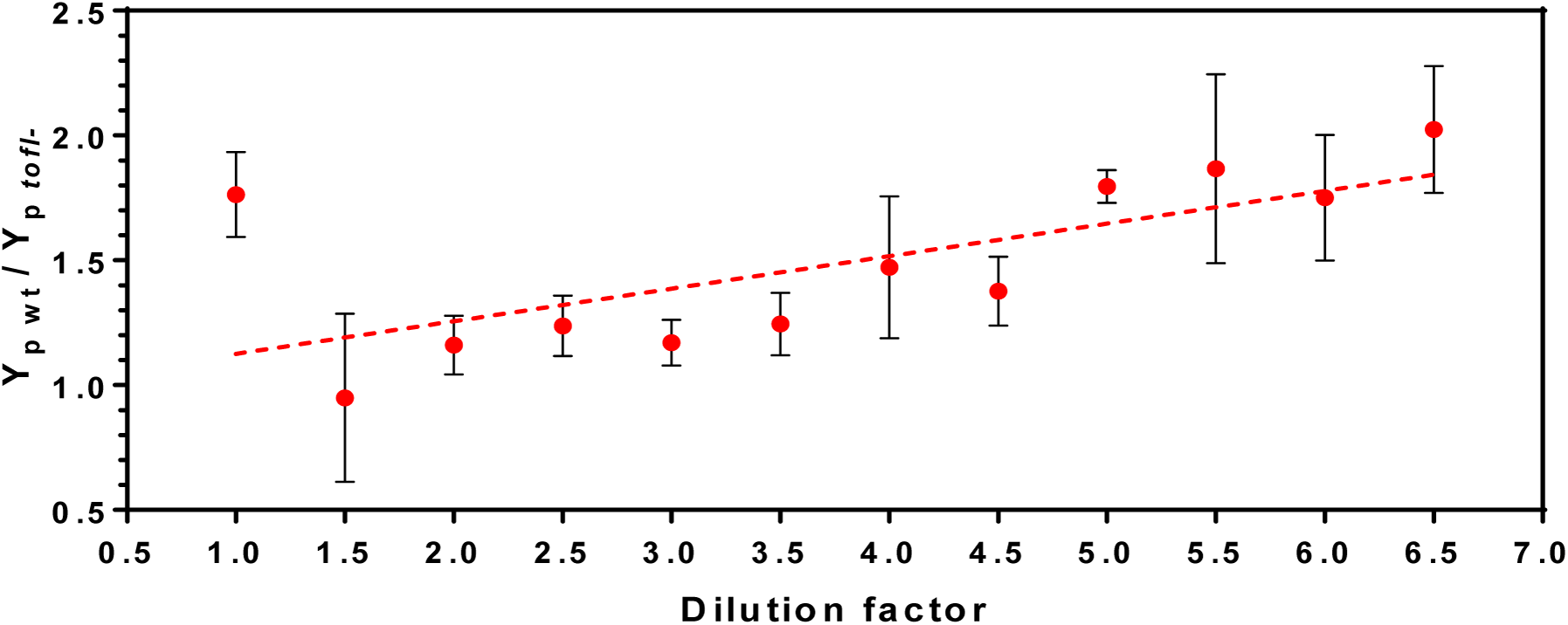
Contribution of quorum sensing in production of rhamnolipids by *B. glumae* in different dilutions of complex NB culture medium. Cultures were performed in increasing dilutions of NB (glycerol maintain stable) medium for 48 h of incubation, then bacterial growth and rhamnolipids production were measured and specific production yield was calculated. The error bars indicate the standard deviation of the mean for three independent experiments.

Diluting the concentration of available nutrients should decrease the growth rate. Thus, to investigate the effect of growth rate on the production of rhamnolipids upon increasing nutrient limitation, the specific growth rate for each NB dilution was determined (Fig. 3). Interestingly, upon decreasing nutrient availability and consequently the specific growth rate (μ), the specific rhamnolipid production yields of the wild-type and *toƒl^-^* mutant were not significantly different until about μ = 0.08 (1/h), mirroring the absence of contribution of QS under unrestricted growth conditions (Fig. 1b). However, from this point on to lower μ values, and in contrast with the QS mutant, we observed a sudden increase in rhamnolipid production yield by the wild-type strain, indicating the initiation of an activating contribution by QS as the growth rate decreases. This reveals a situation where QS-dependent regulation of rhamnolipids is triggered by the growth rate of the population rather than by its cell density.

To better understand this mechanism, we measured the yield of C_8_-HSLs in wild-type strain cultures and uncovered a gradual increase in signal specific concentration as the growth rate decreases (Fig. 3). This means that the number of molecule of C_8_-HSL per unit of cell decreases as the growth rate augments. In addition, we saw that the wild-type C_8_-HSL and rhamnolipid yield curves are never parallel, which further substantiates that additional regulatory elements control the expression of QS-modulated factors such as rhamnolipids.

**Figure 3.**
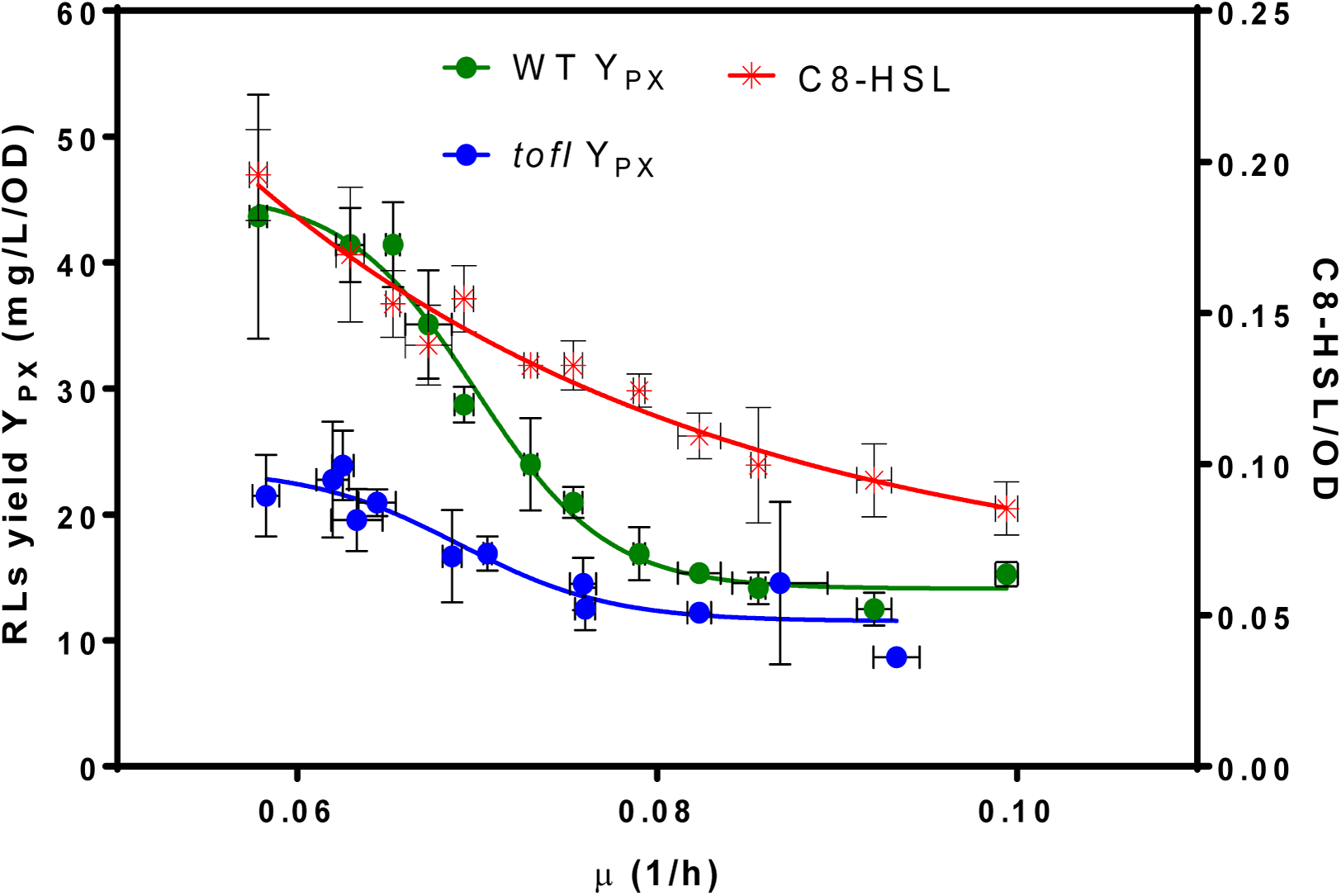
Effect of growth rate on specific yield of rhamnolipids and C_8_-HSL concentration. After 48 h of incubation, rhamnolipids specific production yield in wild-type and *toƒl^-^* mutant and specific yield of C_8_-HSLs in wild-type in different dilutions of NB medium were determined for a range of specific growth rates, determined as described in Materials and methods. The error bars indicate the standard deviation of the mean for three independent experiments.

### Nutritional cues amplify rhamnolipid biosynthesis gene expression through QS-dependent regulation

Finally, to look more closely at the regulatory level where QS is acting, we wanted to know if nutrient-depleted conditions directly influence the expression of QS-dependent rhamnolipid biosynthesis genes. To do so, we monitored *rhlA* transcription in wild-type and *toƒl^-^* strains using a *rhlA*-*lux* reporter gene fusion in different dilutions of NB medium (always with the same glycerol concentration). This allowed us to investigate the direct effect of growth rate on QS-dependent regulation of *rhlA* gene. Hence, we plotted the *rhlA* expression levels in wild-type and *toƒl^-^* strains at a range of specific growth rates. As shown in figure 4, as the growth rate decreases we can observe a sudden increase in *rhlA* gene expression in wild-type strain. However, the expression level of *rhlA* gene in the *toƒl^-^* mutant remains mostly unchanged. Taken together, these data suggests that induction of *rhlA* transcription by TofR occurs at a specific growth rate, instead of specific cell density or AHL concentration.

**Figure 4.**
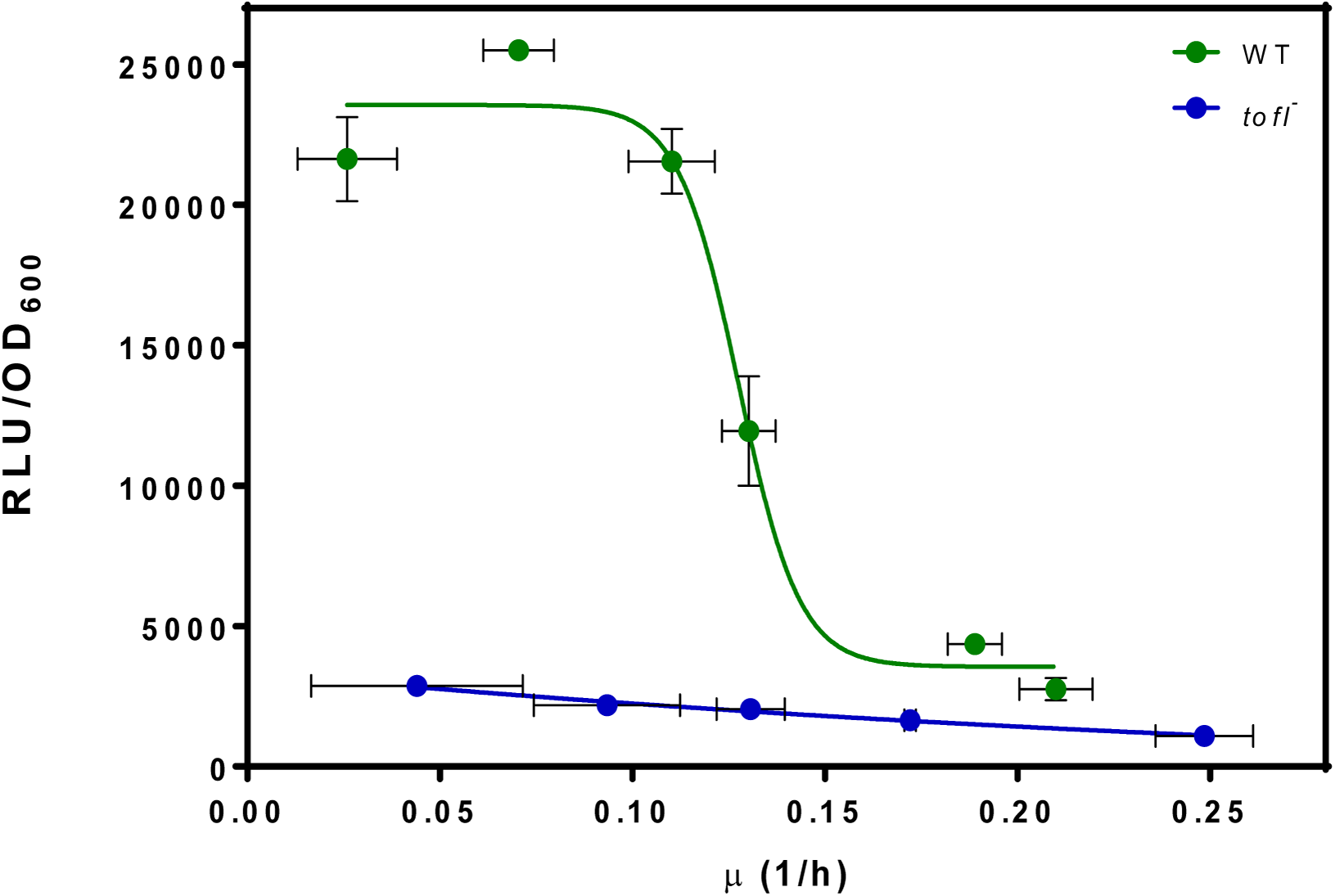
The effect of growth rate on expression from the *rhl* operon promoter in wild-type and *toƒl^-^*strains carrying a chromosomal *rhlA’*-*lux* transcriptional reporter. The *rhlA* promoter activity in wild-type and *toƒl^-^* mutant plotted against growth rates achieved by different dilutions of NB medium. The error bars indicate the standard deviation of the mean for three independent experiments.

## Discussion

We present conclusive evidence for a functional role of QS in providing an optimized demand-driven supply of target cooperative behavior, wherein QS-dependent regulation of rhamnolipids is tightly coupled to specific growth rate. Our results support the proposed model that the adaptive functionality of QS in regulation of public goods is to maintain the homeostasis of cooperation where the production of costly public goods is conditioned by assessment of demand and efficiency (Hense and Schuster, 2015).

We have previously demonstrated that in the rice pathogen *B. glumae*, QS positively regulates the biosynthesis of rhamnolipids, extracellular surface-active glycolipids which are necessary for promoting the social surface behavior called swarming motility (Nickzad et al., 2015). Here, we found that a signal-negative mutant that does not produce C_8_-HSL maintains the ability to produce rhamnolipids in a rich nutrient broth medium, in contrast with growth in a minimal medium where rhamnolipid production is absent (Fig. 1). This led us to hypothesize that QS-dependent regulation of rhamnolipids in *B. glumae* is nutritionally conditional. Comparison of the yield of rhamnolipids per biomass in the wild-type and *toƒl^-^* mutant strains in a decreasing gradient of nutrients revealed that the more we dilute the growth medium the more QS regulation contributes to the production of rhamnolipids (Fig. 2). We increasingly realize that the QS circuitry in various bacteria is highly integrated with other global regulatory pathways, stress responses and specific environmental and physiological cues (Mellbye and Schuster, 2011, Williams and Camara, 2009). Although up-regulation or down-regulation of QS-controlled factors such as rhamnolipids biosynthesis genes *rhlAB* in *P. aeruginosa* by elements other than cell-to-cell communications such as nutritional status is well known (Deziel et al. 2003; Guerra-Santos et al. 1986), this modulation always necessitates the presence of a functional QS system. The identification of a QS-dispensable regulation of rhamnolipids in *B. glumae* adds these metabolites to the list of *B. glumae* factors whose dependence on QS is conditional (Chen et al., 2012, Jang et al., 2014, Kato et al., 2014, Kim et al., 2007).

The dispensability of QS-regulated virulence factors and other costly metabolites, such as rhamnolipids, upon prevailing conditions and environmental cues provides insights into how these regulatory pathways have evolved to optimize the functionality of target genes they regulate. For instance, in the case of rhamnolipids biosynthesis in *B. glumae*, we found that once unrestricted nutrient resources are available, production of rhamnolipids is carried out independently of QS regulation, obviously through still-unidentified regulatory mechanisms. However, under nutrient-limited conditions with excess carbon source, QS becomes the main regulating mechanism significantly enhancing the specific rhamnolipids yield. Under nutrient-limited conditions, which *B. glumae* presumably meets in its natural habitat such as plant tissue environment, it might be more cost-effective for rhamnolipids biosynthesis to be carried out *via* a decision-making mechanism such as QS which places more restrictive controls over nutrient utilization. Notably, QS in *B. glumae* directly controls nutrient acquisition of individual cells in crowded populations (An et al., 2014).

Since growth rate is a direct function of nutrient availability, we hypothesized that growth rate is the key triggering factor that induces QS-dependent regulation of rhamnolipids. Our results indicate there is a growth rate threshold below which QS engages in the regulation of rhamnolipids (Fig. 3). In *P. aeruginosa*, slow growth is one key condition inducing regulation of QS-controlled public good genes in a metabolic prudent manner (Mellbye and Schuster, 2014). For instance, prudent regulation of rhamnolipid production ensures that cells only invest carbon into rhamnolipids biosynthesis when growth is limited by another growth-limiting nutrient, such as nitrogen (Xavier et al., 2011). In fact, presence of excess carbon in different dilutions of NB medium in our experiments, also suggest that in *B. glumae*, QS-dependent regulation of rhamnolipids is coupled to growth rate in a metabolic prudent manner. Furthermore, a gradual increase in QS signal specific concentration upon decrease of specific growth rate suggests a reduction in activation signal threshold, which reflects an increase in cellular demand for production of QS-dependent target gene product at such low density populations. Importantly, our gene expression analyses demonstrate the direct influence of the specific growth rate on QS-dependent induction of *rhlA* expression in the wild-type background, in agreement with a previous report showing that *rhlAB* expression in *P. aeruginosa* in the presence of excess carbon is tightly coupled to the growth rate, and not just cell density (Xavier et al., 2011). Therefore, the role of QS in regulation of public goods is not limited to stationary phase situations at higher cell densities, but also upon encountering restrictive environmental conditions, an early induction of QS by slowed growth rate could provide some inclusive fitness benefits for the survival of an individual cell and could act as a trade-off mechanism between survival and efficiency. Our results are also in accordance with the regulation of QS gene expression by starvation in *P. aeruginosa* in which stringent response can lower the quorum threshold *via* triggering an increased signal synthesis (van Delden et al., 2001, Schuster et al., 2013).

From an ecological significance perspective, induction of QS-dependent regulation of rhamnolipids at low specific growth rates could reflect two distinct scenarios for which such integrated regulation has evolved: (i) restricted nutritional conditions with low cell density (free living bacteria exposed to nutritional stress during establishment of host infection), and (ii) exhausted nutritional environment with a high cell density (bacteria living in biofilms). In both instances, since population growth is limited by the carrying capacity of the direct environment, it is conceivable that slow growth can serve as a predictive factor of the population carrying capacity (de Vargas Roditi et al., 2013). Accordingly, in *B. glumae*, QS is used for anticipation of a population reaching the carrying capacity of an habitat to avoid lethal ammonia-induced alkalization by activating oxalate production, and thus promoting survival in the stationary phase (Goo et al., 2012).

An effective colonization of surface niches necessitates a balance between biofilm and motility (van Ditmarsch et al., 2013). In this regard, there is an inverse regulation of biofilm formation and swarming motility (Caiazza et al., 2007). Since production of surfactants is as crucial for biofilm development as for swarming motility (Nickzad and Déziel, 2014), it seems that the induction of QS-dependent rhamnolipids biosynthesis upon reduced growth rate is a key evolved strategy to modulate swarming motility and promote dissemination of bacterial cells to find new surface niches to colonize (Déziel et al., 2003, Tremblay and Déziel, 2010). Thus, integration of slowed growth rate with QS as a decision-making mechanism for biosynthesis of costly rhamnolipids could serve to assess the demand and timing for expanding the carrying capacity of a population through spatial expansion mechanisms such as swarming motility, thus promoting the chances of survival. In conclusion, we propose that the adaptive significance of growth rate-dependent functionality of QS in rhamnolipids biosynthesis lies within providing a regulatory mechanism for selecting the optimal trade-off between survival and efficiency.

## Materials and methods

### Bacterial strains, plasmids, media and growth conditions

All *B. glumae* strains used are derivatives of the wild-type strain BGR1; the bacterial strains and plasmids used in this study are listed in Table 1. Unless otherwise specified, the strains were routinely regrown from frozen stocks by culturing at 37°C in tryptic soy broth (TSB) (BD) and 240 rpm in a TC-7 roller drum (New Brunswick, Canada), or on TSB agar plates. Antibiotics were used at the following concentrations: tetracycline (Tc), 10 μg ml^-1^, gentamycin (Gm), 20 μg ml^-1^ and triclosan 25 μg ml^-1^.

**Table 1.**
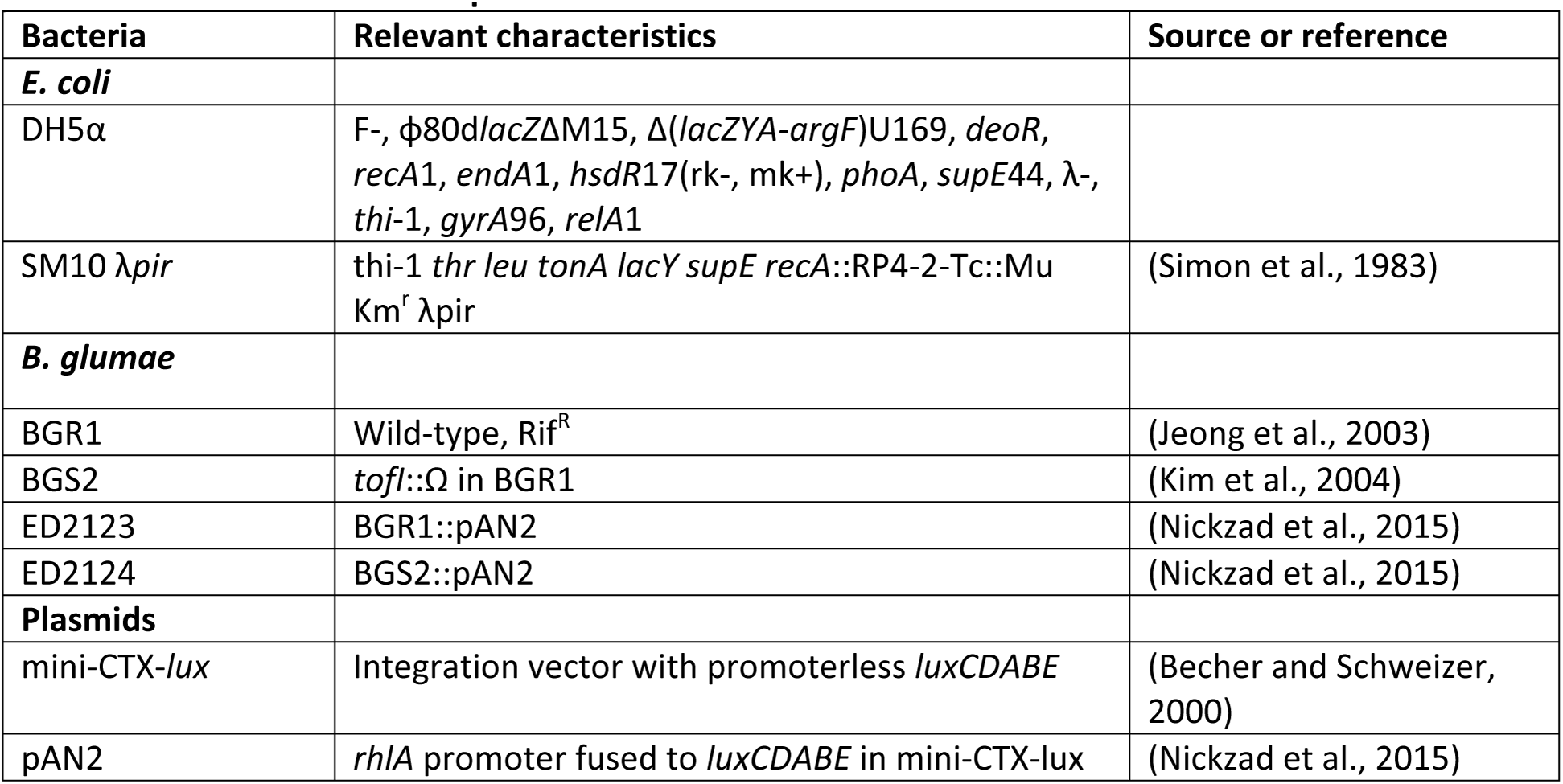
Bacterial strains and plasmids.

For rhamnolipid production, complex nutrient broth (NB) medium (BD Difco, Mississauga, ON, Canada) supplemented with 4% (w/v) glycerol or mineral salt medium supplemented with *2%* (w/v) glycerol and 0.1 M urea was used. The composition of the latter medium was (g/l): KH_2_PO_4_, 2.5; K_2_HPO_4_, 1.5; MgSO_4_, 0.1; CaCl_2_.2H_2_O, 0.1. For NB dilution cultures, glycerol concentration was kept constant at 4% (w/v). To determine whether the rhamnolipid production could be restored by exogenous addition of signal molecules, culture media were supplemented with 8 μM C_8_-HSL. Cultures were grown at 34°C with shaking (240 rpm) in a TC-7 roller drum (New Brunswick, Canada).

### Growth rate measurement

The specific growth rate for wild-type and *toƒl^-^* mutant in each dilution of NB medium was determined using the following equation: 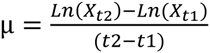 where X_t2_ refers to optical cell density at time t_2_, X_t1_ is the optical cell density at time t1 and μ represents specific growth rate.

### Reporter construction

Transcriptional fusion strains of *B. glumae* were constructed by PCR amplification of a 923-bp fragment containing the promoter region of *rhlA* from BGR1 genome using primers promrhlAF: 5ʹ-ACGTAGAATTCGGGAAAGCAGGCAGGGTAG and promrhlAR: 5ʹ- ACGTAAAGCTTTT TCGATAGGCATGGCGTACTC. The PCR product was digested with *EcoRI* and *HindIII* and ligated within the corresponding sites in mini-CTX-*lux* (Becher and Schweizer, 2000). The resulting construct was integrated into BGR1 and *toƒl^-^* mutant chromosome at the *attB* site through conjugation. Successful chromosomal integration of construct in BGR1 and *toƒl^-^* mutant was confirmed by PCR.

### LC/MS rhamnolipid analysis

The rhamnolipid concentration in the various bacterial cultures was determined by LC/MS (Abdel-Mawgoud et al., 2014). During a period of six days, 400 μl culture samples were retrieved at regular time intervals and the OD_600_ was measured (Nanodrop ND-1000, Thermo Fisher Scientific). Then the samples were centrifuged at 16,000 × *g* for 10 min. to remove the bacteria. To 300 μl of supernatant were added 300 μl of acetonitrile and 10 mg/L 5,6,7,8-tetradeutero-4-hydroxy-2-heptylquinoline (HHQ-d4) as an internal standard (Dubeau et al., 2009). Samples were analyzed by high-performance liquid chromatography (HPLC; Waters 2795, Mississauga, ON, Canada) equipped with a C8 reverse-phase column (Kinetex, Phenomenex) using a water⁄acetonitrile gradient with a constant 2 mmol l-^1^ concentration of ammonium acetate (Dubeau et al., 2009). The detector was a mass spectrometer (Quattro Premier XE, Waters). Analyses were carried out in the negative electrospray ionization (ESI-) mode.

### LC/MS-MS analyses for AHL production

AHL production analysis were carried out by HPLC (Waters 2795, Mississauga, ON, Canada) equipped with a C8 reverse-phase column (Kinetex C8, Phenomenex), and the detector was a mass spectrometer (Quattro Premier XE, Waters). Analyses were carried out in the positive electrospray ionization (ESI+) mode, supplemented by the multiple reactions monitoring (MRM) mode. Samples were prepared as described in (Chapalain et al., 2013).

### Measurement of *rhlA’-lux* activity

Expression from the *rhl* operon promoter in wild-type and QS-negative *toƒl^-^* strains was quantified by measuring the luminescence of cultures of bacteria containing a transcriptional fusion of the *rhlA* promoter with the luciferase gene integrated in the genome. Cells from overnight cultures were washed twice in PBS and inoculated into test medium at an OD_60_0 of 0.05 in white clear bottom 96-well plates (Corning; 3632) containing three replicates of 200 μl per sample. The plates were incubated at 34 °C in a multimode plate reader (Cytation3, BioTek, Winooski, VT) with double orbital shaking. OD_60_0 and luminescence (relative light units, RLU) were measured every 15 minutes for a duration of 48 hr. The luminescence *(rhlA-lux* activity) was adjusted for culture density (RLU/OD_60_0).

## Acknowledgments

Thanks to Professor You-Hee Cho (CHA University, South Korea) for providing strains BGR1 and BGS2. Special thanks to Marie-Christine Groleau for insightful comments and to Sylvain Milot for technical advice. We are grateful to Joao Xavier (Memorial Sloan Kettering Cancer Center, NY) for critical review of the manuscript. Investigations on rhamnolipids in the ED laboratory are funded by the Natural Sciences and Engineering Research Council of Canada (NSERC) Discovery grant No. 312478 and by Team grant 174316 from the Fonds de recherche du Québec-Nature et Technologie (FRQ-NT). AN was recipient of a Ph.D. scholarship awarded by the Fondation Universitaire Armand-Frappier de l’INRS. ED holds the Canada Research Chair in sociomicrobiology.

